# Mechanically Activated Extracellular Vesicle Functionalised Melt Electrowritten Materials for Bone Regeneration: A Mechano-Biomimetic Scaffold

**DOI:** 10.1101/2021.03.29.437528

**Authors:** Kian F. Eichholz, Angelica Federici, Mathieu Riffault, Ian Woods, Olwyn R. Mahon, Lorraine O’Driscoll, David A. Hoey

**Author notes:** **Corresponding Author:** Dr. David Hoey.

## Abstract

Mechanobiological cues arising directly via tissue/scaffold mechanics or indirectly via mechanically activated cell secretomes represent potent stimuli that mediate cell behaviour and tissue adaptation. Exploiting these cues in regeneration strategies holds great promise for tissue repair. In this study, we harness indirect biophysical cues originating from osteocytes in a combination with direct biophysical cues from Melt ElectroWritten (MEW) scaffolds to form a single engineered construct with the aim of synergistically enhancing osteogenesis. The secretome of mechanically activated osteocytes was collected within conditioned media (CM) and extracellular vesicles (EV) were subsequently isolated. Building on MEW micro-fibrous scaffolds with controlled microarchitecture and mineral nanotopography optimised for bone repair, a protocol was developed to functionalise these materials with CM or EVs. Human MSC proliferation was enhanced in both CM and EV functionalised scaffolds. EV functionalised scaffolds were further found to significantly enhance MSC osteogenesis, with enhanced alkaline phosphatase expression, collagen production, and mineralisation compared to control scaffolds. Furthermore, enhanced formation of mineralised nodules was identified in EV functionalised materials. Combining direct biophysical cues provided by the fibrous architecture/mineral nanotopography with the indirect cues provided by EVs, these constructs hold great promise to enhance the repair of damaged bone in a physiologically relevant manner.

## 2 Introduction

There are many cases where bone is unable to self-repair following damage to the tissue, including large defects and fractures following trauma or tumour growth [1]. The clinical challenges presented by this are significant, and there are no satisfactory solutions to date. Approximately 2,000,000 bone grafting procedures are undertaken each year [2], with the majority of these utilising autografts. However, this approach to bone repair has many issues, resulting in an unmet need for the development of alternative tissue regeneration strategies. Synthetic tissue engineering approaches are an attractive solution and may yield a potentially limitless tissue source for unrestricted repair. Of fundamental consideration when developing this type of approach is the understanding of the cell’s native environment, which is pivotal for mediating its behaviour and ensuring continuous tissue regeneration in addition to guiding repair. In bone, this environment is comprised of an intricate network of several cell types within a complex composite matrix, which collectively co-ordinate cell behaviour in response to the dynamic biochemical and biophysical environment [3]. A fundamental component within this system is the stem cell niche [4] where two regions of particular interest are the endosteal and periosteal surfaces, each of which house specific stem cell populations known to contribute to bone formation and repair [5, 6]. These populations are tightly regulated directly by the endo/periosteal fibrous environment, and indirectly by biological factors released by adjacent cells. Understanding and implementing an environment which mimics the native biochemical and biophysical cues of the stem cell niche would provide a powerful strategy to expand and direct stem cells to guide tissue repair.

A potent regulator of stem cell behaviour and bone regeneration is physical loading. Direct biophysical regulation of stem cells occurs via the underlying fibrous micro architecture and mineral nano-topography of bone tissue. *In vivo*, this micro architecture is composed of fibres with diameters of 1 – 10 μm [3], at varying degrees of alignment depending on local tissue strain [7]. Furthermore, it is known that substrate architecture [8] and stiffness [9] can have profound effects on cell behaviour, in turn influencing cell mechanosignaling and driving stem cell lineage commitment [10]. We have recently demonstrated that stem cell shape and mechanosignaling is significantly altered in response to fibrous architecture specifically [11]. We have shown that a 90° fibre architecture with fibres of 10 μm diameter significantly enhances stem cell osteogenesis over more aligned or random scaffolds, which ultimately occurs as a result of enhanced cell spreading and cytoskeletal tension being imposed by this architecture. A large proportion of bone is composed of inorganic mineral which is closely integrated with the collagen structure to form a composite matrix. The basic mineral unit consists of nano-scale needles with approximate dimensions of base 5 nm and length 50 – 100 nm [12]. These needles form platelets composed of partly merging crystals of the same base and diameter, with platelet widths of 20 – 30 nm, which in turn form stacks and aggregates with irregular 3D structures of size 200 – 300 nm. The scale and architecture of this nano structure is known to be important for stem cell proliferation and differentiation, with stem cell behaviour being negatively affected when cultured on larger particles [13]. In addition, we have recently demonstrated that fibres coated with nano-needle hydroxyapatite (nnHA) can have a profound influence on stem cell osteogenesis [14]. Remarkably, cell mineralisation is enhanced over 14-fold on nnHA fibres compared to uncoated controls, and almost 3-fold greater than fibres coated with micron-scale hydroxyapatite with a plate architecture. In addition, this nano-topography facilitates enhanced protein adsorption and controlled release, mimicking the excellent protein stability provided by the needle-shaped mineral structure of native bone [15]. Thus, implementing defined nano-topographical mineral features is a promising strategy to form biomimetic scaffolds and further enhance osteogenesis.

Indirect biophysical regulation also provides important information to recruit, expand and mediate stem cell behaviour. In bone, paracrine signalling coordinated by the osteocyte is a key component to this effect, with the release of mechanically induced factors driving the behaviour of osteoblasts, osteoclasts and their progenitors [16]. The important role of osteocyte mechanosignaling on stem cell behaviour in particular has been well documented, with conditioned medium from mechanically stimulated osteocytes enhancing the recruitment, proliferation and osteogenic differentiation of stem cells [17, 18]. This is achieved via the release of a range of factors, with several of particular interest including sclerostin [19] and RANKL [20]. We have demonstrated via proteomic analyses that a large range of factors are released by osteocytes, many of which are mechanically regulated, and can enhance stem/stromal cell osteogenesis [21]. It has become apparent that the means by which these factors are delivered is also of great importance. Extracellular Vesicles (EVs) are membrane bound particles released by cells which are implicated in cell signalling in many tissues throughout the body including bone. Bone cells, including osteocytes [22–24], osteoblasts [25–27] and stem cells [28] have been shown to release EVs, which efficiently target cells to deliver signalling components including proteins and miRNAs. In addition to the above, we have also demonstrated that the mechanical conditioning of osteocytes via fluid shear generates mechanically activated osteocyte derived extracellular vesicles (MA-EVs), which significantly enhance stem cell recruitment and osteogenesis over EVs secreted by statically cultured cells. [21]. This further reinforces the key role of osteocyte EVs in indirect biophysical regulation of stem cells and indicates the great promise of osteocyte derived EVs in the development of therapeutics and strategies for bone regeneration. Previous studies have explored the use of EVs for these purposes, which may be loaded with drugs for use as a delivery vector to treat osteoporosis [29], or incorporated within scaffolds to enhance bone regeneration [30, 31]. The further use of MA-EVs for scaffold functionalisation holds great promise, where they may be used to locally guide cell behaviour and enhance regeneration.

In this study, we aim to combine both direct and indirect biophysical cues within the bone stem cell niche to create a mechano-biomimetic bone tissue regeneration strategy. We utilise a previously developed, fibrous 90° scaffold architecture fabricated via melt electrowriting (MEW), as a foundation upon which to build our scaffold. This framework is then coated with nano needle hydroxyapatite (nnHA), which closely recapitulates the natural nano topography of bone to further enhance osteogenesis. In addition, osteocytes are mechanically stimulated and either conditioned medium or subsequently isolated mechanically activated EVs (MA-EVs) are used to further functionalise scaffolds to provide cues for indirect biophysical regulation (Figure 1A). Human bone marrow stem/stromal cells (hMSCs) were then cultured on these scaffolds and their osteogenic capacity investigated via various means, including alkaline phosphatase (ALP) activity, collagen production and mineralisation. We demonstrate that MA-EV functionalised scaffolds significantly enhance stem cell osteogenesis and highlight the synergistic potential of direct and indirect biophysical cues of the bone stem cell niche in driving regeneration.

**Figure 1.**
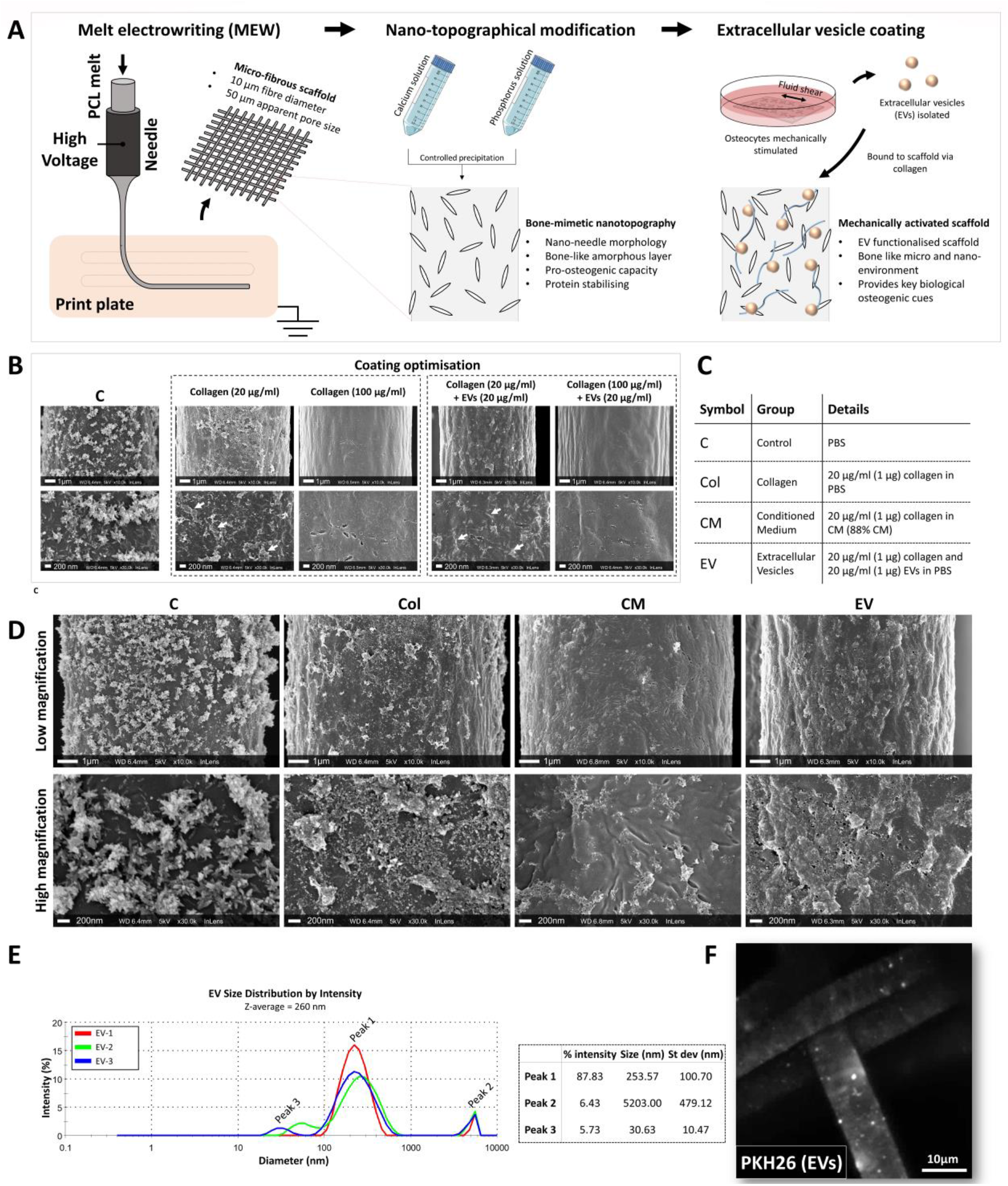
**A** Development of mechano-biomimetic scaffold via MEW, nano-hydroxyapatite coating and EV functionalisation. **B** Development of scaffold coatings, where collagen concentration was optimised for the collagen and EV groups. Mineral needles could still be seen in the lower concentration groups (white arrows), but were almost completely covered in the high concentration groups. **C** Final treatments and concentrations used. **D** SEM imaging of final groups. **E** Particle size distribution in EV group, which consists of a primary peak at approximately 200 nm, and two minor peaks above and below this (n=3). **F** PKH26 staining of functionalised scaffolds demonstrating successful EV binding to scaffolds.

## 3 Results

### 3.1 Fabrication of Conditioned Media (CM) and Extracellular Vesicle (EV) functionalised scaffolds

The effect of collagen coating concentration after 1 h incubation on fibre topography was investigated via SEM imaging (Figure 1B). At a concentration of 100 μg/ml, the collagen almost completely covers the nnHA layer, with some needles partially visible through the collagen coating. At a concentration of 20 μg/ml, a thinner collagen coating is achieved with the nnHA layer largely visible. The influence of the above collagen concentrations in addition to 20 μg/ml of EVs on fibre topography was also investigated. As previously demonstrated, the higher concentration results in a relatively thick collagen/EV coating which masks the needle topography of the nnHA layer. The lower concentration in addition to 20 μg/ml EVs yields a coating which partially covers the nnHA layer with needle topography largely maintained. The effect of coating time was also investigated, where 1 h and 24 h were compared, and was found to have no effect indicating that the coating process happens quickly. The lower collagen concentration at 1 h treatment was thus used for all study groups with the exception of the control (Figure 1C), with SEM images of all groups shown in (Figure 1D).

### 3.2 Characterisation of CM and EVs

Osteocyte derived CM and isolated EV samples were characterised by particle size analysis (n=3) and successful binding to scaffolds was investigated via fluorescent imaging. The primary peak in CM particle distribution, at an intensity of 60.33%, revealed an average particle size of 186.67 nm which is within the range typical of EVs (Figure S 1Figure 1A). A second peak with an intensity of 20% and average particle size of 0.72 nm is likely an experimental artifact, and is too low for the readings typically attained for protein size via DLS [32, 33]. This may be caused by the presence of phenol red or other components within the DMEM culture medium in the CM group. As the sample was filtered prior to analysis via DLS, peak 3 is a likely contaminant. The primary peak in the EV group has an intensity of 87.83% and average particle size of 253.57 nm, which is also within the typical range for EVs (Figure 1E). As previously, the peak below this is a possible artifact which is only present in 2 out of the 3 replicates, while the larger particle peak is a likely contaminant. Functionalised scaffolds were then stained with PKH26, a membrane dye used as a marker to confirm the presence of EVs on EV coated scaffolds. A consistent spread of high intensity point staining throughout the scaffold confirming successful binding of EVs (Figure 1F). Background staining was detected in C and Col groups. In CM there was evidence of EV staining, however, at a relatively low intensity and concentration (Figure S 1B).

### 3.3 Attachment and proliferation of hMSCs

Seeding efficiency is not significantly altered between groups, which have an average cellular attachment of 46% after 24 h (n=5) (Figure 2A). A greater difference in cell number between groups becomes apparent at D7, with DNA content lowest in C at this time and 2.7 fold greater in EV. This effect becomes more apparent at D14, with all groups significantly greater than C by more than 2 fold. Interestingly, at D21 there was a reduction in the amount of DNA detected in Col, CM and EV groups, while DNA content increased in C. This is likely due to the development of a highly dense matrix after 21 days in culture, with constructs being noticeably much stiffer when handling compared to those at earlier time points. This made DNA extraction more difficult, with cells/matrix still partially visible on scaffolds after vigorous vortexing.

**Figure 2.**
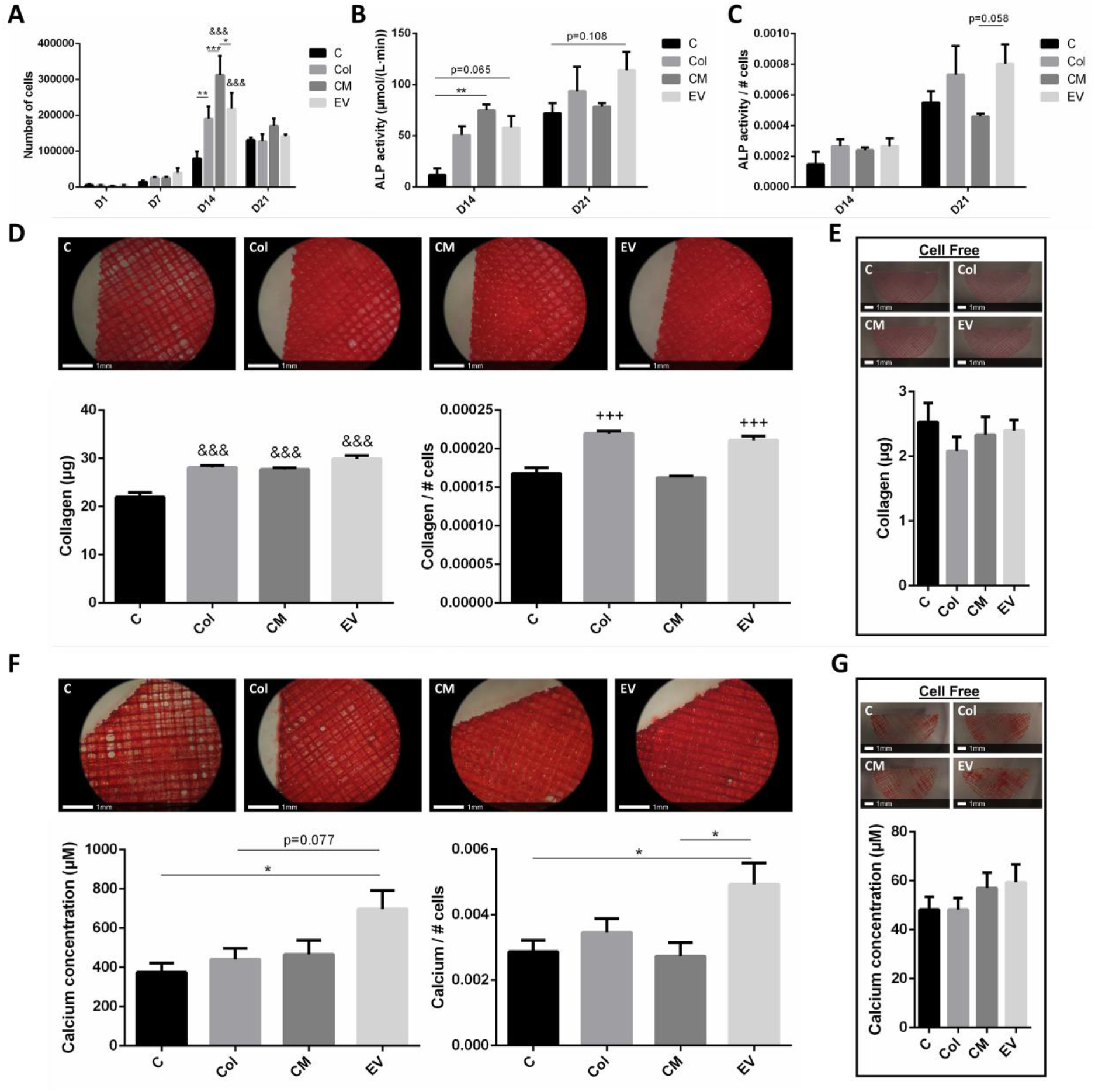
**A** Proliferation quantified via DNA content (n = 5). (& = statistical significance compared to C). **B** Total ALP activity in hMSCs at D14 and D21. **C** ALP normalised to cell number (n=5). **D** Collagen production at D21, where a significant increase of Col and EV compared to C and CM is seen when collagen is normalised to cell number (n=5). **E** Collagen was quantified in cell-free scaffolds after 21D culture, with only mild background staining identified (n=5), confirming that all collagen in cell-scaffold constructs is produced by the MSCs. **F** Cell mineralisation as measured by calcium content at D21 (n=5), where both total calcium content and calcium normalised to cell number is significantly enhanced in the EV functionalised group compared to the control. **G** Calcium was quantified in cell-free scaffolds with a trend towards greatest levels in CM and EV groups (n=4-5).

### 3.4 MSC osteogenesis

#### 3.4.1 ALP activity

Total ALP activity was investigated at D14 and D21 (n=5) (Figure 2B). At D14, ALP is enhanced in the Col group compared to C, however this is not significant. The addition of mechanically activated osteocyte CM for scaffold functionalisation further enhances this effect, resulting in a significant 6.3-fold increase in ALP compared to C. ALP is also enhanced in the mechanically activated EV (MA-EV) group, with a near-significant 4.9-fold increase vs C. At D21 the differences between groups are attenuated, with ALP activity greatest in the EV group at this time with a 1.6-fold increase vs C. ALP activity normalised to DNA was also investigated (Figure 2C). As previously demonstrated, values are lowest in C after 14 days. At D21, ALP/DNA is greatest in the EV group, with a 1.8-fold change vs CM. In summary, the functionalisation of scaffolds with collagen enhances ALP activity compared to control scaffolds, however, the further functionalisation of scaffolds with CM or EVs greatly enhances these effects, indicating the potential of these biological components for further driving stem cell differentiation in fibrous micro-environments.

#### 3.4.2 Collagen production

To further assess the osteogenic capacity of scaffolds, the degree of collagen deposition was assessed (n=5). It can be visually seen that C has a less well developed collagen matrix than all other groups (Figure 2D). Quantifying total collagen reveals that all groups have at least 25% more collagen than C, with the greatest increases in the EV group with 36% more collagen at a total value of 30 μg. Normalising to DNA content, both EV and Col groups have significantly more collagen compared to C and CM. In order to assess the contribution of collagen originally used for scaffold coating towards this result, cell-free scaffolds were cultured in the same conditions up to 21 days (Figure 2E) (n=5). There was visually no difference between groups which appeared to have a mild pink background stain, with this being confirmed upon further quantification. There are no differences between groups, including the control which originally contained no collagen. It can therefore be deduced that none of the original collagen coating remained after 21 days in culture. In conclusion, collagen coating of scaffolds enhances the deposition and stabilisation of collagen matrix by cells, with only this cell-synthesised collagen remaining after 21 days.

#### 3.4.3 Mineral production

Mineralisation was also assessed at D21 as a late-stage marker of osteogenic differentiation (n=5). As previously seen with collagen, visual differences could be seen between scaffold groups (Figure 2F). In particular, C was covered in noticeably less mineralised matrix and reduced staining intensity. In addition, the EV group appeared to have the most intense staining, with a dense, consistent matrix completely covering the scaffolds. Scaffolds in the EV group were also noticeably stiffer when handling. Upon quantification, it was confirmed that C had the lowest level of mineral at 376 μM. A marginal increase of 66 μM was seen in the Col group, with a further increase of 25 μM in CM. A further substantial increase of 231 μM was seen in the EV group compared to CM. This corresponds to a significant 1.9-fold increase in mineralisation in the EV group compared to C, and a 1.6 fold increase vs Col. Similar trends are seen when normalising results to DNA, with significant increases in mineralisation in the EV group compared to C and CM. While all groups are coated with the same quantity of mineral prior to cell culture, cell-free scaffolds were cultured to D21 (Figure 2G) (n=5) to investigate our hypothesis that EVs may act as sites of mineral nucleation in scaffolds, as has been previously demonstrated in 2D culture [25]. While differences in total mineral are marginal, there is evidence that EVs, which are present in both the CM and EV groups, may act as sites of mineral nucleation in fibrous scaffolds independent of cells, as there is a trend towards increased mineral content in these groups after 21 days. In summary, MA-EV functionalised scaffolds significantly enhance mineralisation after 21 days in culture, revealing their potential for driving stem cell osteogenic differentiation and bone formation in fibrous 3D environments.

#### 3.4.4 Mineral characterisation

SEM imaging and EDX analysis was conducted on C and EV groups at day 21 to further characterise the engineered tissue, as these are the least and most osteogenic groups respectively. No differences were identified at low magnifications via SEM imaging, with both groups being completely covered in matrix (Figure 3A), and no observable difference in cellular organisation under high voltage imaging (Figure 3B). At higher magnifications, fibres were almost completely covered in cells, with spherical nodules present both on and adjacent to fibres (Figure 3C). In mid pore regions, greater numbers of these nodules were identified in the EV group (Figure 3D). EDX analyses were conducted to further investigate the mineral structure of the C and EV groups, with minimal differences between them at low magnifications (Figure S 2). To further investigate the structure of nodules which are present in both groups, EDX mapping was done on clusters of nodules at high magnification. Calcium and phosphorus were found to have a greater intensity within these clusters (Figure 3E), indicating their potential role as sites of mineral nucleation. Ca/P ratios were investigated in cell-free constructs, with a value of 1.75 in C and a significantly greater Ca/P ratio of 1.91 the EV group (Figure 3F). This is likely due to the presence of minerals in EVs which may influence the mineral composition of the nnHA coating. Interestingly however, Ca/P ratios in cell-scaffold constructs were much lower, with values of 0.84 and 0.70 in C and EV groups respectively (Figure 3G). These values were recorded in mid-pore regions to investigate cell-produced mineral while limiting the influence from the nnHA coating on fibres.

**Figure 3.**
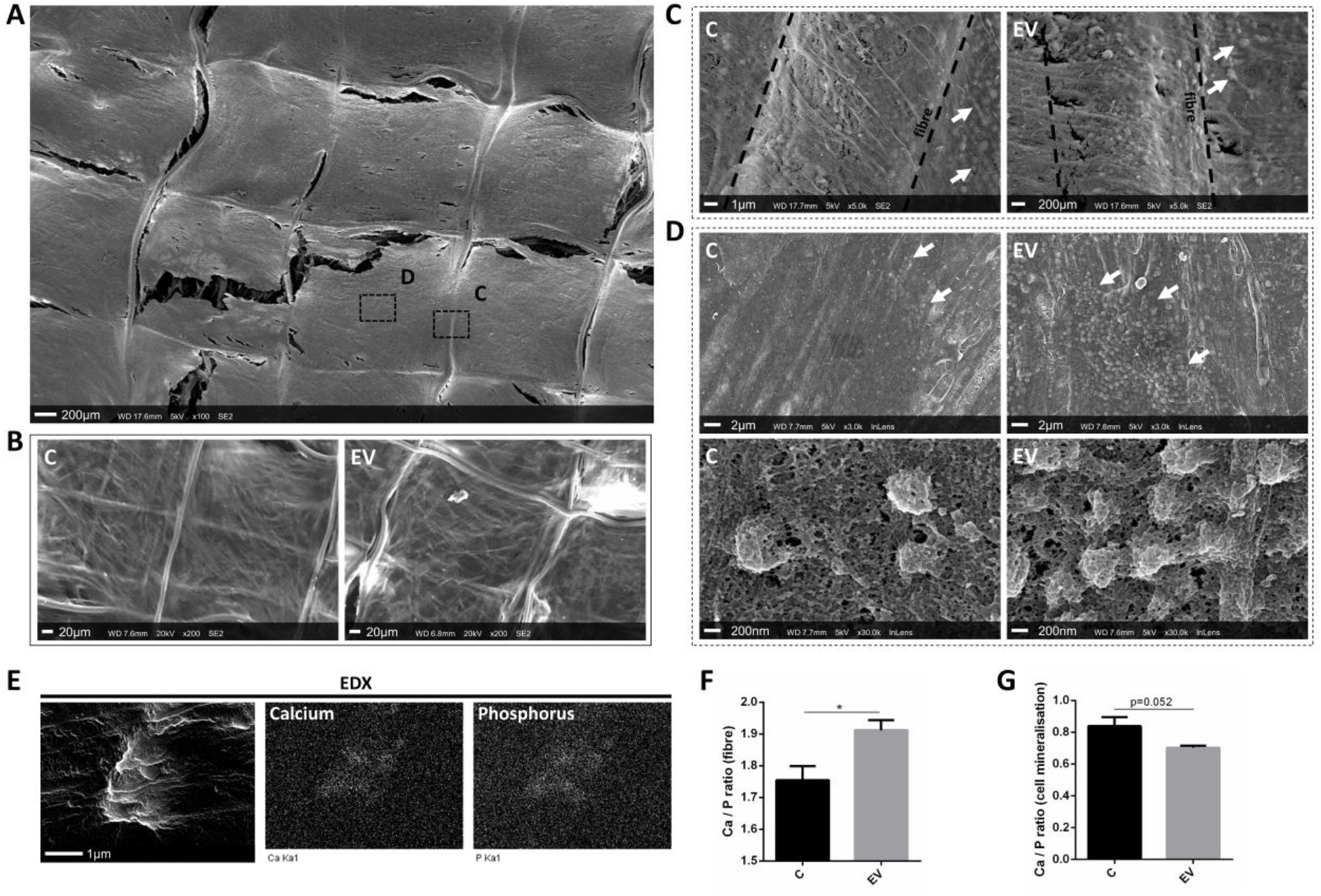
SEM imaging of C and EV constructs at D21. **A** Both groups were enveloped in matrix where two regions of interest, C and D are highlighted. **B** High voltage imaging revealing the underlying cellular organisation. **C** Superficial fibres are almost completely covered, and the nHA coating can be seen through the matrix in addition to the presence of spherical nodules (white arrows). **D** Spherical nodules may also be seen in the mid-pore regions, and are more numerous in the EV group. **E** Element mapping was conducted on a cluster of nodules, where the greater intensity of calcium and phosphorus in this region confirms it as a site of mineral nucleation. **F** Ca/P ratios were investigated in fibres of cell free constructs at D21 (n=5) and **G** were also investigated in cell-scaffold constructs at D21 (n=5).

## 4 Discussion

Mechanobiological cues represent potent stimuli that mediate cell behaviour and coordinate tissue adaptation. Exploiting these cues by incorporation into regeneration strategies holds great promise for the repair of many tissues throughout the body. In this study, we detail the development of a fibrous scaffold combining direct and indirect biophysical cues which enhance stem cell osteogenesis to effectively guide bone tissue regeneration. We show that components, namely mechanically activated osteocyte derived conditioned medium and isolated EVs, when functionalised onto MEW scaffolds can enhance stem cell proliferation and osteogenic differentiation as seen by elevated ALP expression, collagen deposition, and mineralisation. This approach elicited a greater response to that previously shown with BMP2 functionalisation [14]. This is further verified via SEM imaging and EDX analysis, where increased formation of mineralised nodules was seen in EV functionalised scaffolds. These findings demonstrate that indirect biophysical cues provide a powerful strategy for enhancing stem cell proliferation and mineralisation within 3D environments. Combined with the direct physical cues provided by the scaffold architecture and topography, this mechano-biomimetic approach has great potential to guide local bone tissue regeneration and repair.

Many strategies for bone tissue regeneration are comprised of engineered matrices which may be supplemented with potent factors such as bone morphogenic proteins (BMPs). While this has widely been demonstrated to significantly enhance *in vitro* osteogenesis and *in vivo* defect healing, there has recently been many concerns raised over the long-term safety of this approach. Adverse effects including osteolysis, ectopic bone formation and increased cancer risk have all been reported following *in vivo* use of BMPs [34–36]. While these factors are naturally occurring within bone, the local imbalance caused by the high concentrations of an individual factor utilised in an engineered scaffold is likely the leading contributory factor to the adverse effects reported above [37]. This highlights the need for an alternative approach to stimulate bone repair in a more physiologically relevant manner. It is widely known that mechanical loading enhances the size and mineral content of bone [38], and exploiting the natural means by which this occurs holds great promise for development of tissue regeneration strategies. It is now known that the osteocyte plays a key role in this process, sensing mechanical loads and communicating to other bone residing cells via secreted factors to mediate their behaviour [17, 18, 39, 40]. We have further demonstrated that EVs are a primary means by which osteocytes deliver these factors, with mechanical stimulation of osteocytes enhancing the recruitment and differentiation capacity of released EVs [21]. Furthermore, the great stability of the lipid membrane of EVs in addition to their specificity when delivering cargo to target cells [41] makes them a prime candidate for the delivery of regenerative factors via engineered scaffolds to enhance bone repair in a safe and physiologically relevant manner.

One of the challenges of utilising EVs within engineering scaffolds entails securely binding them to the underlying structure. The majority of previous approaches have achieved this via loading an EV solution directly on to the scaffold before incubation to allow binding [31, 42–46]. One consideration with this method is the efficiency of EV binding. There is considerable work involved in the production of conditioned medium and isolation of EVs, and this loading method can result in a significant loss of these components depending on the scaffold material. Another method which has been reported entails suspending EVs within a gelatin solution before forming a hydrogel with EVs encapsulated inside [47], which significantly reduces any loss of EVs. We have functionalised our scaffolds with a combinatory approach incorporating elements of both the above methods, with the foundation being a nHA coated scaffold. This mineral coating has nano-topographical features and a high surface area which stabilises proteins and enhances binding [48]. In addition to the enhanced binding properties provided by the nHA mineral, we have further re-suspended CM and isolated EVs within a collagen solution, which is subsequently applied to scaffolds to further enhance binding efficiency. Membrane staining validated the presence of EVs on the fibre surface, both in CM, and to a greater extent, EV functionalised scaffolds. An important consideration in this work was to preserve the equally important direct biophysical cues provided by the micro-fibrous matrix and nano-topographical features of the mineral coating, both of which play important roles in driving stem cell osteogenesis [11, 49]. Collagen concentration was thus optimised to preserve these features and provide a thin CM/EV enriched coating to further enhance these regenerative effects via mechanically activated biologics. This is the first study which has incorporated EVs with a defined micro-fibrous matrix to yield a scaffold which drives cellular behaviour via both direct and indirect biophysical cues.

The long-term influence of CM and EV functionalised scaffolds on proliferation and osteogenesis was investigated by culturing scaffolds with hMSCs. Consistent with previous findings which demonstrate that osteocyte CM enhances MSC proliferation [50], we have demonstrated that CM, and to a lesser extent, EV functionalised scaffolds exhibit a similar effect. Furthermore, we have demonstrated through several markers the great capacity of EVs in driving MSC osteogenesis on fibrous scaffolds. Previous studies have reported on the capacity for EVs in promoting bone regeneration via immobilisation on scaffolds [31, 43–45]. EVs from mechanically stimulated osteocytes have also been shown to enhance bone formation [24]. However, this is the first report of osteocyte EVs being used for scaffold functionalisation, with the additional mechanical activation step providing significant benefits by exploiting the highly mechanosensitive nature of osteocytes to enhance mineralisation. Of particular interest is the means by which EVs enhance regeneration, and in particular, mineralisation. EVs are rich in annexin proteins and membrane bound lipids which enhance calcification [51]. They have been shown to act as sites for mineral nucleation, and remarkably, to significantly enhance calcium deposition compared to BMP2 [25]. We have shown that mineralised nodules, closely resembling those previously identified in mineralising osteoblasts [52, 53], are present in both C and EV groups in close proximity to nnHA coated fibres. In mid pore regions, however, there are a greater number of nodules in the EV group indicating likely cellular uptake of EVs to globally enhance mineralisation. Also of particular interest is the Ca/P ratio of cells at day 21, which ranges between 0.70 – 0.84. This closely matches the value of 0.75 previously found for intracellular mineral-containing vesicles [54] while also corresponding closely to the Ca/P ratio of amorphous calcium phosphate (ACP) [55, 56], a precursor phase which later transforms within collagen fibrils to hydroxyapatite [54]. Further from the significantly enhanced mineralisation in MSC seeded EV scaffolds, we have also seen marginal increases in mineral content in cell-free EV scaffolds after 21 days, in addition to elevated Ca/P ratios of the fibre mineral coating. This provides evidence that EVs alone may mineralise in culture. Taken together, it is evident that EVs play an important role in mineralisation and enhance further mineral deposition by MSCs cultured on EV functionalised scaffolds. It is thus clear that EVs hold great promise not only as a means to recruit cells and drive behaviour via delivery of numerous signalling factors, but also as sources of mineralisation, with promising applications for the functionalisation of scaffolds to guide bone regeneration in a biomimetic manner.

## 5 Conclusions

In summary, we have developed for the first time a scaffold which incorporates elements of direct biophysical stem cell regulation via a defined fibrous and mineral architecture, along with indirect biophysical regulation provided by mechanically activated EVs from stimulated osteocytes. The capacity for EVs to enhance osteogenesis, and particularly mineralisation is clear, and this approach thus has great potential for applications in bone regeneration. In addition, this biomimetic approach overcomes the issues associated with using highly potent factors such as BMPs and has the potential to facilitate guided regeneration in a safe and physiologically appropriate manner.

## 6 Experimental Section

### 6.1 Melt electrowriting (MEW)

Fibrous scaffolds with a fibre diameter of 10 μm, square apparent pore size of 50 μm, and layer fibre spacing of 300 μm were fabricated on a custom built MEW apparatus as previously described [11]. Briefly, heated air at a temperature of 90°C was circulated around a syringe to melt polycaprolactone (PCL) (Sigma Aldrich 440744, average Mn 80,000), with air pressure being used to extrude the polymer through a 21G needle with high voltage applied at a distance of 15 mm from a grounded aluminium collector plate. Fibres were deposited with a 90° offset between subsequent layers to result in a square pore shape.

### 6.2 Nano-needle hydroxyapatite (nnHA) coating

A calcium solution was made with 0.05M calcium chloride dihydrate (Sigma Aldrich C7902) in MilliQ water. A phosphate solution was made with 0.03M sodium phosphate tribasic dodecahydrate (Sigma Aldrich S7778) and 0.01M sodium hydroxide in MilliQ water. Reactions were carried out in 50ml conical tubes, with reagent volumes being maintained at 40 ml at each step of the process and 6 scaffolds of dimension 3 x 3 cm being processed per tube. Scaffolds were immersed in 70% ethanol for 15 min under vacuum. They were then placed in 2M NaOH, vacuum applied for 5 min and incubated at 37°C for 45 min. Scaffolds were then rinsed 5 times in MilliQ water and added to 20 ml of the calcium solution. 20 ml of the phosphate solution was slowly added to the calcium solution, vacuum applied for 5 min and samples incubated in the calcium/phosphate solution for 30 min at 37°C. This coating procedure was repeated a further two times minus the vacuum treatment. Scaffolds were then treated with 0.5M NaOH for 30 min at 37°C, rinsed 5 times in MilliQ water and allowed to dry overnight.

### 6.3 Osteocyte cell culture

MLO-Y4 osteocyte like cells (Kerafast) [57] were maintained as previously described [58] in α-MEM growth medium with 2.5% fetal bovine serum (FBS), 2.5% calf serum (CS), 1% Penicillin/Streptomycin (PS) and 1% L-glutamine. 6-well plates were coated with 0.15mg/ml type I collagen (Sigma C3867) for one hour and washed with PBS, after which osteocytes were seeded at a density of 1.16 x 10^4^ cells/cm^2^ in 2 ml medium/well. After 48 h culture, plates were then placed on an orbital shaker for 2 h at a rotational speed of 100 rpm. This mechanical stimulation regime yields an average fluid shear stress of 0.28 Pa and maximum force of approximately 1 Pa across the bottom of the well [59]. Cells were washed with PBS and 833 μl of serum free medium (1% PS and 1% L-glutamine) was applied to each well. Cells were incubated for 24 h and conditioned medium (CM) was then collected. Samples were centrifuged at 3,000g for 10 mins at 4°C to remove debris, after which the supernatant was collected.

### 6.4 Extracellular vesicle isolation from conditioned media

Medium was filtered through a 0.45 μm pore filter and ultracentrifuged (Beckman Coulter Optima KPN-100) at 110,000 g for 75 min at 4°C, using an SW32.Ti swing bucket rotor. Pellets were washed in cold particle-free PBS and the ultracentrifugation process was repeated. EVs were then re-suspended in particle-free PBS and protein quantity determined via a NanoDrop spectrophotometer with absorbance of 280 nm. CM and EV particle size distribution was investigated via digital light scattering (DLS) on a Zetasizer Nano ZS. Samples were first diluted 1:50 in PBS and filtered using a 0.45 μm filter.

### 6.5 Functionalising MEW constructs with EVs and CM

Scaffolds were punched to a diameter of 8 mm and UV sterilised for 20 min on each side before being placed in 48-well plates. Scaffolds were then pre-wet in a graded ethanol series of 100%, 90% and 70% for 20 min each before being washed three times in sterile MilliQ water and allowed to dry overnight. Scaffolds were functionalised with EVs or CM within a collagen matrix to aid adhesion to fibres, and variables for collagen coating were optimised to obtain a coating which preserves the nano-topography of the mineral coating. The influence of time (1 hr and 24 hr) on collagen coating thickness was investigated at a collagen concentration of 100 μg/ml. The influence of collagen concentration (20 μg/ml and 100 μg/ml) on collagen coating thickness after 1 hr treatment was investigated. Final scaffold treatment methods are as follows. For the EV group, a 50 μl solution in PBS containing 1 μg collagen (20 μg/ml) and 1 μg EVs (20 μg/ml) was loaded on each scaffold. For the CM groups, a 50 μl solution of 88% CM containing 1 μg collagen (20 μg/ml) was loaded on to each scaffold. Control (C) scaffolds were loaded with PBS, and collagen control (Col) scaffolds were loaded with 1 μg collagen. Scaffolds with above treatments were incubated for 1 h at 4°C and were then rinsed in PBS.

### 6.6 SEM imaging and Energy-dispersive X-ray spectroscopy

For scanning electron microscope (SEM) imaging, samples were prepared for imaging by coating with gold/palladium for 40 s at a current of 40 mA. For analysis with energy-dispersive X-ray spectroscopy (EDX), scaffolds were carbon coated and analysed at a voltage of 15kV in a Zeiss ULTRA plus SEM with an 80mm^2^ Oxford Inca EDX detector. To investigate approximate calcium/phosphorus atomic ratios for each group, spectra were acquired on scaffolds for 120 s (n=5 technical replicates). Element maps were constructed at a resolution of 512 x 384 with map dwell of 4000 μs and linescan dwell of 2000 μs.

### 6.7 Immunofluorescent staining of MEW-EV constructs

Fluorescent membrane labelling was used to confirm the presence of EVs on functionalised scaffolds. Constructs were incubated with 1 μM PKH26 dye solution (PKH26GL, Sigma) for 5 min. Excess dye was then quenched by treatment with EV depleted FBS (3 x 5 min treatments). Constructs were then washed in PBS (3 x 5 min) and mounted to glass slides using Fluoroshield (Sigma Aldrich, F6182) mounting medium. Fluorescent imaging was carried out using a Leica SP8 scanning confocal microscope with 20x objective, where Z-stacks with 10 steps and a total thickness of 10 μm were constructed.

### 6.8 Human Bone Marrow Mesenchymal Stem/Stromal (hMSC) Cell culture

Human bone marrow mesenchymal stem/stromal cells (hMSCs) were isolated from bone marrow (Lonza, US), trilineage potential verified (data not shown), and seeded at a number of 10,000 cells per scaffold. Scaffolds were transferred to new well plates after 24 h, and cultured in 300 μl osteogenic medium (OM)/well from day 3, which consisted of 10% FBS DMEM supplemented with 100 nM dexamethasone, 10 mM β-glycerol phosphate and 50 μg/ml ascorbic acid. Medium was changed every 3.5 days.

### 6.9 Proliferation

At days 1,7,14 and 21, scaffolds were added to 100 μl lysis buffer in 1.5 ml tubes (n=5) containing 0.2% Triton X-100, 1 mM Tris pH8, with phenylmethylsulfonyl fluoride (PMSF) being added at a ratio of 1:200 just before use, and stored at −80°C. Before DNA quantification, samples were subjected to three freeze-thaw cycles in liquid nitrogen and underwent sonication for 60 s. Samples were then vigorously vortexed before being stored on ice for processing. DNA content was quantified using a Quant-iT™ PicoGreen™ dsDNA Assay Kit (Invitrogen, P7589), with excitation and emission wavelengths of 485 nm and 528 nm respectively. The DNA content in 10,000 hMSCs seeded and pelleted in centrifuge tubes was also quantified to calculate the total number of cells present on scaffolds.

### 6.10 Characterisation of hMSC osteogenic differentiation

#### 6.10.1 Intracellular ALP

Intracellular ALP was quantified at days 14 and 21 (n=4). 50 μl of 5mM pNPP was added to wells along with 10 μl of cell lysate and 70μl MilliQ water in a 96-well plate. Standards were comprised of serial dilutions of p-Nitrophenyl phosphate (pNPP, Sigma Aldrich, N1891) with 10μl of 43μM ALP enzyme (Sigma Aldrich, P6774) being added to each. Plates were incubated for 1 h in the dark at room temperature, and reactions were then stopped using 20μl of 3M NaOH. Absorbance was read on plates at 405 nm, and ALP activity was calculated as the amount of pNPP generated as a function of sample volume and reaction time.

#### 6.10.2 Collagen production

Scaffolds were cultured up to 21 days before evaluating collagen content. Cell-scaffold constructs were rinsed in PBS, fixed in 10% neutral buffered formalin for 15 min and rinsed again in PBS before storage at −20°C. Scaffolds were stained with 200μl of 1mg/ml of Direct Red 80 (Sigma Aldrich, 365548) in a saturated aqueous picric acid solution for 1 h with shaking at 150 rpm. Scaffolds were then washed twice with 0.5% acetic acid and allowed to dry overnight before imaging. To quantify collagen content, 500μl 0.5M NaOH was added to wells under shaking until stain was dissolved, and solutions were added to 1.5 ml tubes. Tubes were centrifuged at 14,000g for 10 min to pellet debris. Standards were made by adding direct red staining solution to 8μl of collagen I (Corning, #354249) before centrifuging at 14,000g for 10 min and re-suspending the collagen in 500μl 0.5M NaOH. The absorbance of samples and standards were read at 490 nm in 96-well plates.

#### 6.10.3 Calcium production

Cell-scaffold constructs were investigated for total calcium content after 21 days. Cell-free scaffolds were also cultured up to 21 days to determine the contribution of total calcium from cell mineralisation versus mineral nucleation due to the presence of EVs. Scaffolds were incubated with 1% alizarin red S solution (Sigma Aldrich, A5533) in distilled water for 10 min at a pH of between 4.1 – 4.3. Samples were rinsed 3 times with water and allowed to dry prior to imaging. To quantify bound stain, 400 μl of 10% acetic acid was applied and samples incubated at room temperature for 30 min while shaking at 150rpm. The acetic acid was added to 1.5 ml tubes, vortexed vigorously and heated to 85°C for 10 min. Tubes were transferred to ice for 5 min, centrifuged at 20,000g for 15 min, and 300 μl of the supernatant was added to new tubes along with 120 μl of 10% ammonium hydroxide. Standards were made with dilutions of alizarin red solution in distilled water, with the pH of each adjusted to between 4.1 – 4.3. Samples and standards were read at 405 nm in a 96-well plate.

### 6.11 Statistical analysis

Subsequent biological data is presented in terms of average and standard error of the mean. For proliferation and ALP data, statistical analysis was performed using two-way ANOVA and Tukey’s multiple comparison post-test. For collagen and calcium data, statistical analysis was performed using one-way ANOVA and Tukey’s multiple comparison post-test. For EDX data, statistical analysis was performed using unpaired Student’s t-test.

## 7 Supplementary information

**Figure S 1.**
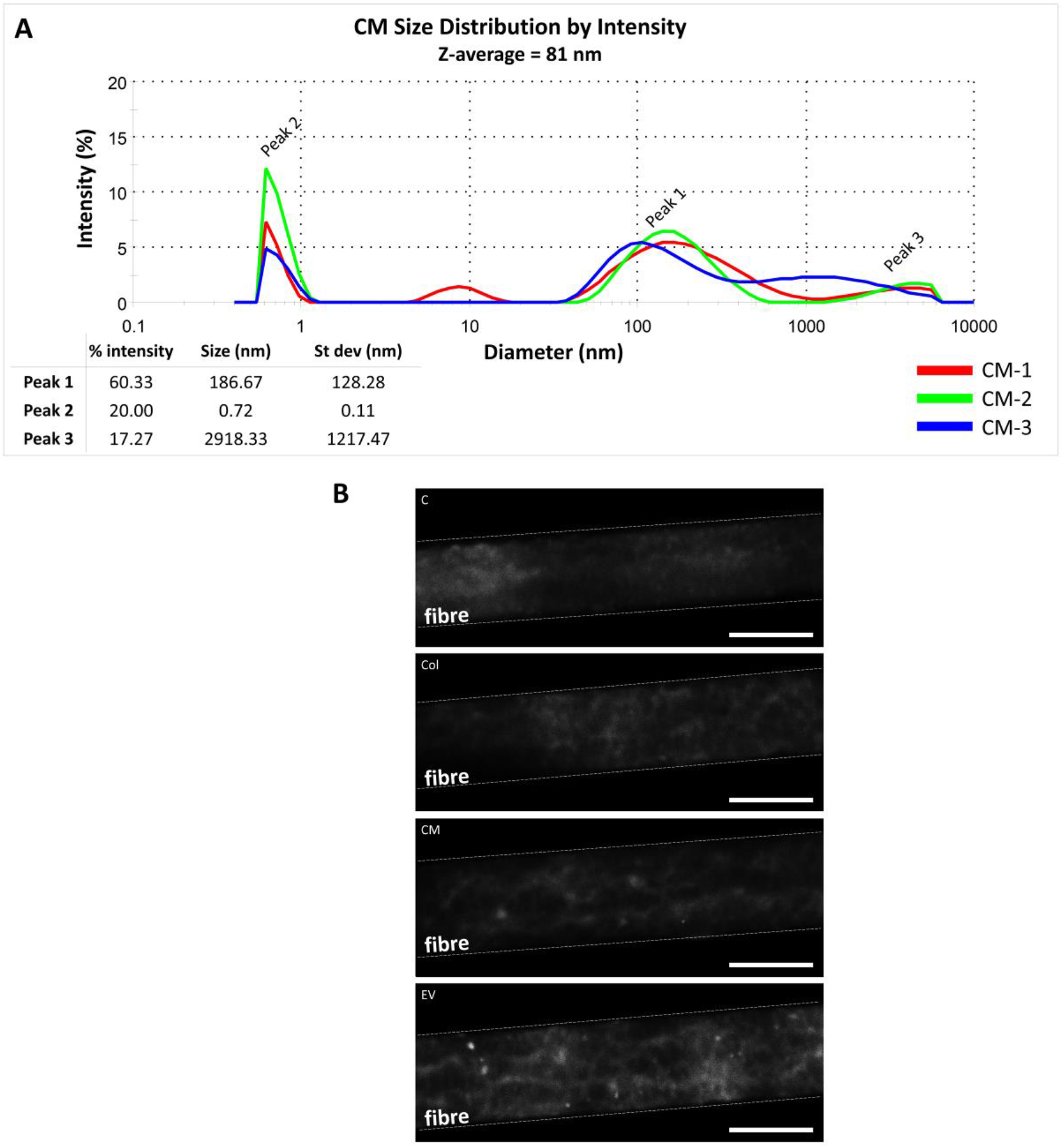
**A** Particle size distribution in CM group, which consists of a primary peak at approximately 200 nm, and two minor peaks above and below this. **B** PKH26 staining of functionalised scaffolds demonstrating successful EV binding in the CM and EV groups.

**Figure S 2.**
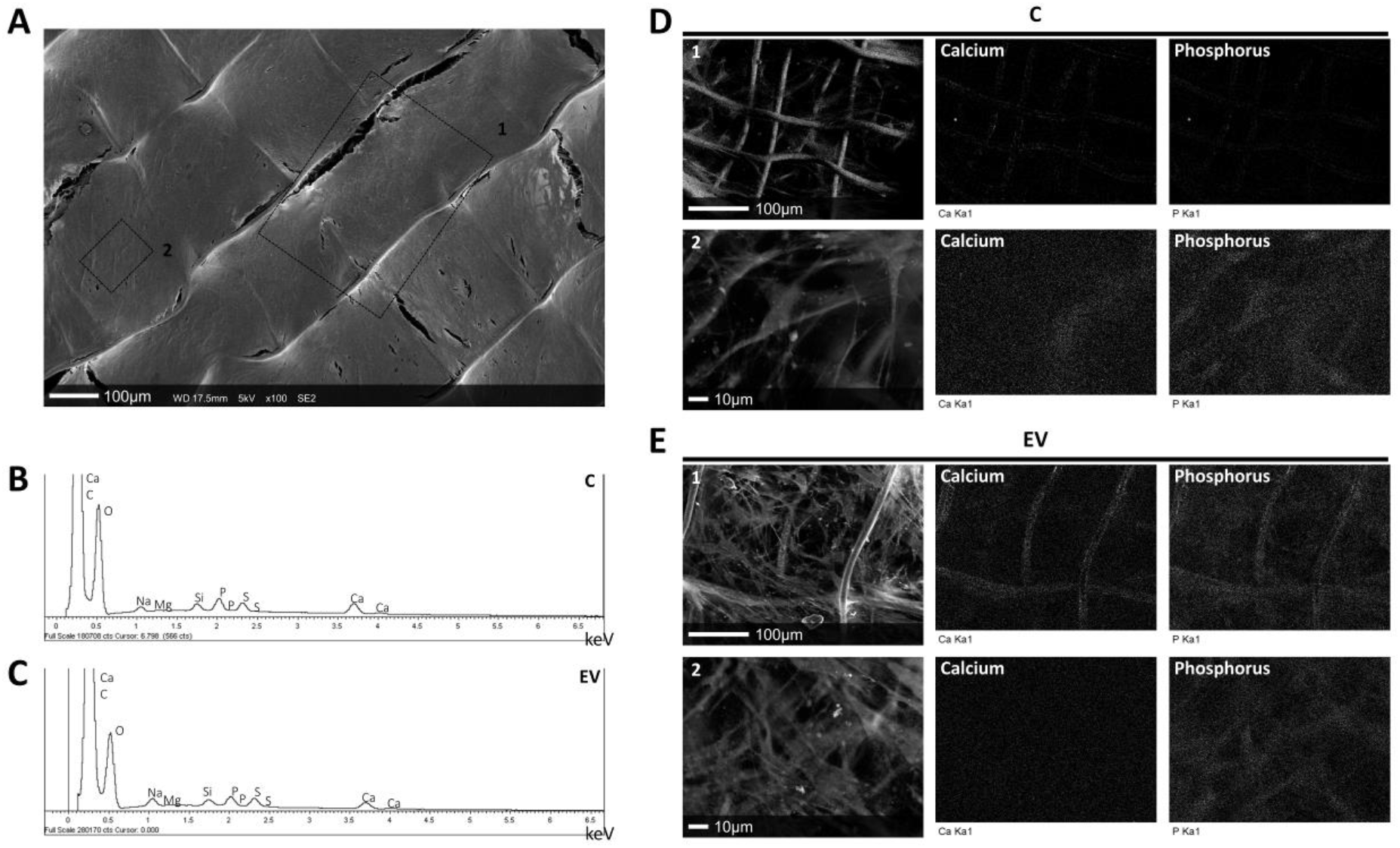
EDX analysis of constructs at D21. **A** Element mapping was conducted in regions 1 and 2 for C and EV scaffolds, as these groups have the lowest and highest osteogenic capacity respectively at D21. **B-C** Spectra were similar for both C and EV groups. **D-E** Element maps show a consistent spread of calcium and phosphorus in both groups.

